# Transcriptomic Signatures for Ovulation in Vertebrates

**DOI:** 10.1101/069716

**Authors:** Dongteng Liu, Michael S. Brewer, Shixi Chen, Wanshu Hong, Yong Zhu

## Abstract

Recently, we found anovulation in nuclear progestin receptor (Pgr) knockout (Pgr-KO) zebrafish, which offers a new model for examining Pgr regulated genes and pathways that are important for ovulation and fertility. In this study, we examined expression of all transcripts using RNA-Seq in pre-ovulatory follicular cells collected after the final oocyte maturation, but prior to ovulation, from wild-type (WT) or Pgr-KO fish. Differential expression analysis revealed 2,888 genes significantly differentially expressed between WT and Pgr-KO fish. Among those, 1,230 gene transcripts were significantly more expressed, while 1,658 genes were significantly less expressed in WT than those in Pgr-KO. We then retrieved and compared transcriptional data from online databases and further identified 661 conserved genes in fish, mice, and humans, that showed similar levels of high (283 genes) or low (387) expression in animals that were ovulating compared to those with no ovulation. For the first time, ovulatory genes and their involved biological processes and pathways were also visualized using Enrichment Map and Cytoscape. Intriguingly, enrichment analysis indicated the genes with higher expression were involved in multiple ovulatory pathways and processes such as inflammatory response, angiogenesis, cytokine production, cell migration, chemotaxis, MAPK, focal adhesion, and cytoskeleton reorganization. In contrast, the genes with lower expression were mainly involved DNA replication, DNA repair, DNA methylation, RNA processing, telomere maintenance, spindle assembling, nuclear acid transport, catabolic processes, nuclear and cell division. Our results indicate that a large set of genes (>3,000) are differentially regulated in the follicular cells in zebrafish prior to ovulation, terminating programs including growth and proliferation, and beginning processes including the inflammatory response and apoptosis. Further studies are required to establish relationships among these genes and an ovulatory circuit in zebrafish model.

## Introduction

Ovulation is a physiological process that releases a fertilizable oocyte from follicular cells and is an essential reproductive event for the preservation of a species. It is well established that luteinizing hormone (LH) initiates a cascade of signaling; including upregulation of progestin and its nuclear progestin receptor (PGR) which activates various downstream targets and signaling pathways, eventually leading to follicular rupture. However, our understanding of the molecular mechanisms that control ovulation is far from complete. For example, there is limited evidence of downstream targets and signaling pathways that PGR regulates. A few genome-wide transcriptome analyses of differentially expressed genes in the follicular cells of pre-ovulatory oocytes suggest conserved gene expression regulation in humans, macaques, and mice [1–3]. To our knowledge, there are no published transcriptomic analyses of gene expression focusing on the follicular cells of pre-ovulatory oocytes in basal vertebrate; all studies conducted so far have not separated follicular cells from the oocytes, or used mixed stages of oocytes that preclude detailed comparisons between different species [4–6].

Zebrafish is an alternative vertebrate model for studying gene function, signaling pathways in development, and various physiological processes because of their low cost, rapid development, and relative simplicity. Unlike mammalian models, zebrafish release and fertilize mature oocytes outside the body, developing their embryos externally. Because embryos develop externally, females do not undergo cumulus-oocytes complex (COC) expansion, luteinization, or implantation processes that happen concurrently or subsequently with ovulation, making it relatively easy to distinguish genes exclusively involved in ovulation [7]. In addition, follicular cell layers can be collected separately from pre-ovulatory oocytes, which are relative large (>600 µm), for biochemical and molecular analyses [8]. Our recent study has also shown that Pgr is an important transcription factor induced by luteinizing hormone (LH), and is essential for ovulation in zebrafish [9]. In Pgr knockout (Pgr-KO) female zebrafish mature oocytes are trapped within follicular cells unable to ovulate, leading to infertility. Our results are consistent with the complete anovulatory and infertile phenotype reported in PGR-KO mice [9, 10]. These results prompted us to hypothesize that ovulation is controlled by conserved genes and signaling pathways in vertebrates.

In this study, we first conducted a genome-wide differential gene expression analysis in the follicular cells of pre-ovulatory oocytes from wildtype (WT) in comparison to Pgr-KO zebrafish using RNA-Seq and bioinformatics tools. We hypothesize that gene expression changes in WT would be important for ovulation, while lack of changes in Pgr-KO due to anovulation could serve as reference. We then conducted a comparison analysis of genome-wide differentially regulated genes in the follicular cells of pre-ovulatory oocytes of three key vertebrate species, i.e., zebrafish, mouse, and human. We found that Pgr regulates a network of conserved signaling pathways, biological processes, and genes that control ovulation. Dramatic differences in the expression between WT and Pgr-KO of various genes include *ptgs2* (prostaglandin-endoperoxide synthase 2a, 2b), *runx1* (runt-related transcription factor 1), *ptger4b* (prostaglandin E receptor 4b), *rgs2* (regulator of G-protein signaling 2), and *adamts9* (a disintegrin-like and metalloproteinase with thrombospondin motifs). This differential expression across species provides a list of candidate genes to study the molecular mechanisms underlying ovulation.

## Materials and Methods

### Zebrafish husbandry

Generation and characterization of Pgr mutant lines have been described previously [9]. The WT zebrafish used in this study was a Tübingen strain initially obtained from the Zebrafish International Resource Center and propagated in our lab according to the following procedure. Fish were kept at constant water temperature (28^0^C), a photoperiod of 14hrs of light with 10 hrs of dark (lights on 9:00, lights off at 23:00), pH at 7.2, and salinity conductivity from 500-1200 μS in automatically controlled zebrafish rearing systems (Aquatic Habitats Z-Hab Duo systems, Florida, USA). Fish were fed three times daily to satiation with a commercial food (Otohime B2, Reed Mariculture, CA, USA) containing high protein content and supplemented with newly hatched artemia (Brine Shrimp Direct, Utah, USA). The Institutional Animal Care and Use Committee (IACUC) at East Carolina University approved experimental protocols.

### Collection of pre-ovulatory follicular cell layers

Follicular cells of pre-ovulatory oocytes were collected from three WT or three Pgr-KO female zebrafish at the same developmental stage, i.e., immediate after oocyte maturation but prior to ovulation (See Fig. 1 for detail). We limited our sample size to n = 3 for each group, to balance high cost of RNA-seq and minimum requirement of statistical analyses. Follicular cells are two thin layers of cells (<4 µm in thickness) containing theca and granulosa cells, surrounding a gigantic oocyte in zebrafish (Ø>600 µm for pre-ovulatory oocyte, Fig. 1). Follicular cells could be physically separated from oocytes [8], which typically have approximate 1000 fold higher amount of total RNAs than those in surrounding follicular cells. However, physical separation of granulosa cells from theca cells is impractical due to small cell size and no distinguishable physical properties of these two cell types.

**Figure 1.**
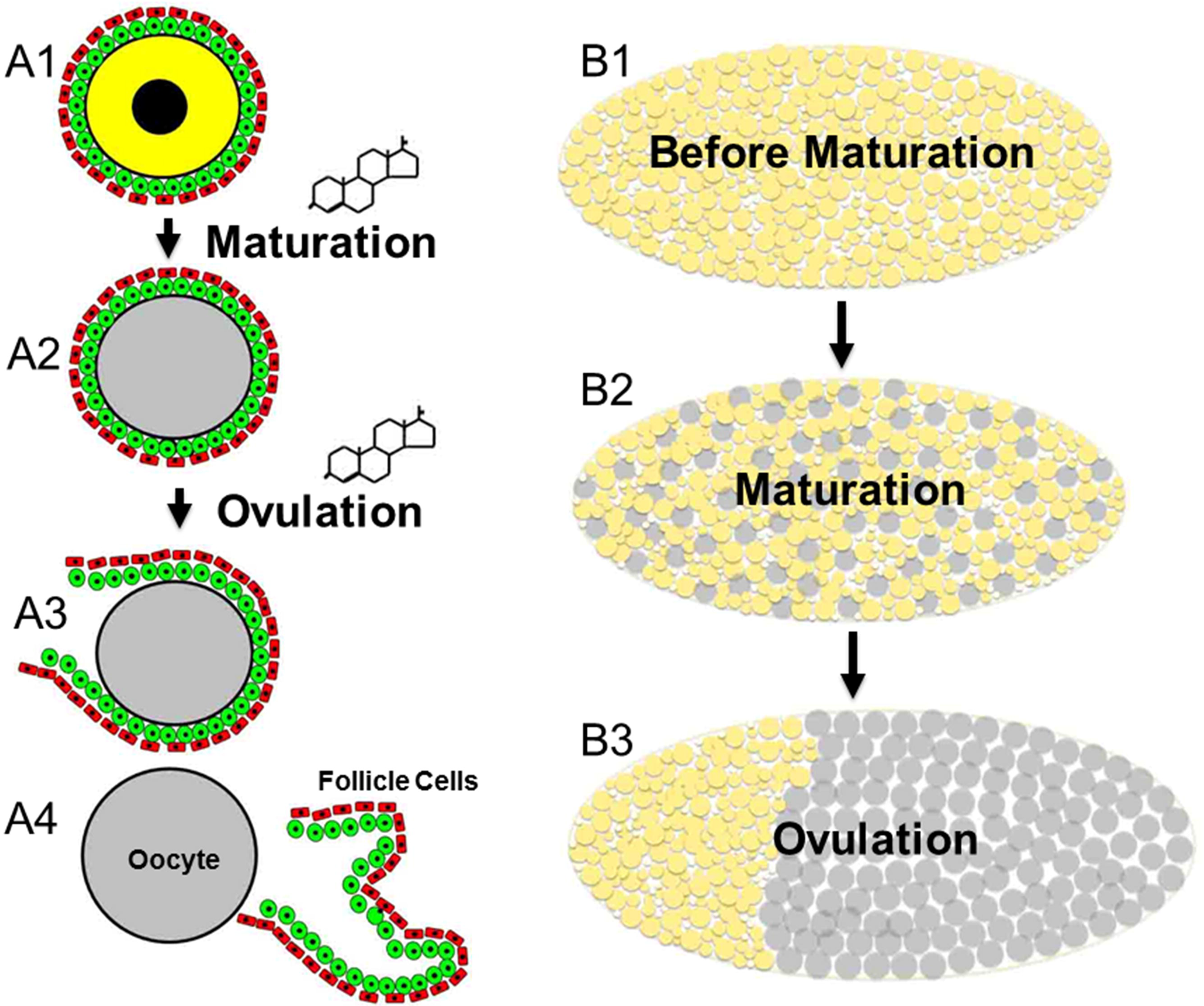
Final oocyte maturation (FOM) and ovulation are easily observable and distinguishable with naked eyes in zebrafish. Representative maturing oocytes are on the left (A1-A4), and representative changes in ovaries are on the right (B1-B3). FOM (resumption of meiosis) occurs prior to ovulation. The maturation process includes germinal vesicle break down (GVBD), chromosome condensation, assembly of meiotic spindle, and formation of the first polar body. Change in cytoplasm appearance of stage IV oocytes from opaque (in yellow) to transparent (in light gray) is the most reliable indication that oocyte maturation is complete (from A1 to A2, or B1 or B2) in zebrafish. Once ovulation is complete, these mature and transparent oocytes migrate to the posterior of animal body (A2 to A3, and A4; or B2 to B3), ready to be released and fertilized outside. These recurring tissue remodeling processes, including maturation and ovulation, happen daily in wildtype (WT) zebrafish. In contrast, oocyte growth and maturation occur normally but ovulation was completely blocked in Pgr knockout (Pgr-KO). Matured oocytes were absorbed before next wave of oocytes going through GVBD and FOM in Pgr-KO females. Follicular cells used in this study were all collected from same stage of oocytes, i.e. after maturation but prior to the ovulation (panels A2 & B2) from WT or Pgr-KO fish.

All the fish used in the experiment were approximately four months old. Individual, well-fed, mature, and healthy female zebrafish were housed separately from male fish by a middle divider in a spawning tank the night before sampling. Oocyte maturation and ovulation were synchronized between different individuals, and spawning typically happened within 30 minutes once the lights switched on and the middle divider was removed from these well-fed and individually housed fish. To obtain pre-ovulatory oocytes, ovaries were removed within an hour before the lights turned on in the morning following an appropriate anesthetic overdose (MS-222: 200 mg/L in buffered solution). Excised individual ovary was then placed in a zebrafish Ringer’s solution (116mM NaCl, 2.9mM KCl, 1.8 mM CaCl2, 5mM HEPES, pH 7.2) and examined under a dissecting microscope. Ovaries containing healthy pre-ovulatory follicles that had a translucent appearance, indicating the completion of oocyte maturation and proximal to ovulation, were selected (Fig. 1). Ovaries with no pre-ovulatory oocytes, incomplete absorption of previous left-over matured oocytes, or unhealthy oocytes that were morphologically distinct from healthy pre-ovulatory oocytes were discard. Individual, follicle-enclosed, healthy mature oocytes were teased away from immature oocytes by gently pipetting in and out several times using a Pasteur pipette. To determine cell specific changes of transcripts regulated by Pgr in the follicular cells, we manually peeled follicular cells off pre-ovulatory oocytes in zebrafish Ringers’ solution under a dissecting microscope using a pair of fine, clean glass needles as described previously [8]. Follicular cells were collected into a 1.7ml microcentrifuge tube and homogenized immediately in a 300 µl TRIzol solution by a sonicator (Sonic Dismembrator, Fisher Scientific, Pittsburgh, PA, USA). Samples were stored in –80 ℃ freezer until RNA extraction. Each sample contained follicular cell layers separated carefully from 45 to 130 pre-ovulatory oocytes of one fish. The process for collecting one sample was limited to less than an hour, avoiding significant degradation or changes of the transcripts (unpublished data). We collected six samples in six different days, in order to sample at same time point and same developmental stages.

### RNA isolation

Total RNA was extracted using TRIzol and a Qiagen RNeasy kit according to a modified protocol. An equal volume of cold 100% ethanol was added into the aqueous phase of the solution following phase separation TRIzol. Samples were then loaded onto an RNeasy spin column, centrifuged (8,000*g*, 30 seconds), washed once with 700 μl RW1, twice with 500 μl RPE washing buffer, and eluted in 25 μl of RNase free water according to Qiagen’s instructions. The approximate concentration and purity of samples were examined using a Nanodrop 2000 Spectrophotometer. An aliquot of RNA sample with OD 260/280 > 1.8 and OD 260/230 > 1.6 was used for RNA-Seq analysis and the remainder was used for quantitative real-time PCR (qPCR) assay.

### Library construction and Illumina sequencing

RNA-Seq library preparation and high-throughput NGS sequencing was carried out at HudsonAlpha Genomic Services Lab (Huntsville, AL). Qubit® 3.0 Fluorometer and Agilent 2100 Bioanalyzer further examined the concentration and integrity of total RNA samples prior to library construction. About 800ng of total RNA was used to construct a cDNA library according to a protocol of Illumina TruSeq RNA Sample Preparation Kit (Illumina). Ribosomal reduction was used to remove non-coding rRNA. The library was then PCR amplified with 15 cycles using TruSeq indexes adaptor primers, submitted for Kapa quantification and dilution, and sequenced with a single end read (50bp) on an Illumina HiSeq 2000 instrument.

### Genome mapping and differential expression analysis

The quality control and adaptor trimming of raw FastQ files were performed using Trim_Galore. Trimmed raw files were inspected using FastQC. The entire zebrafish genome sequence (version GRCz10) was downloaded from Ensembl, and the alignment of sequence reads to the zebrafish genome was carried out using STAR aligner [11]. Binning of sequencing reads to genes/exons was accomplished by HTseq-count, where reads with an alignment score of less than 10 were skipped [12]. DESeq2, a popular Bioconductor package with over 3000 citations, was chosen to normalize the raw counts with respect to the gene length and sequencing depth as well as identify differentially expressed genes [13].

### Validation of differential expression by quantitative real-time PCR (qPCR)

Twenty-three of the differentially expressed genes were selected, and their expressions were further validated using traditional qPCR. Briefly, 250 ng of total RNA from a subset of samples that were used for transcriptomic analysis, were reverse transcribed using SuperScript III Reverse Transcriptase in a 10 μl reaction volume following the manufacturer’s instructions (Invitrogen, Carlsbad, CA). Specific PCR primer pairs (supplemental Table S1) for target genes were designed as close as possible to stop codon to detect transcripts without five-prime caps, and spanning at least two adjacent exons to avoid genomic DNA interference. Glyceraldehyde-3-phosphate dehydrogenase (*gapdhs*) was chosen as an internal control for qPCR because *gapdhs* was expressed evenly among all samples in RNA-Seq analysis. Absolute copy numbers of each transcript were calculated from a standard curve generated from a serial dilution of plasmid DNA with known concentrations [8]. Then, the expression of each transcript was normalized with *gapdhs* and expressed as fold changes by comparing WT to knockout (log2 (WT/Pgr-KO)), since genes important for ovulation would change significantly in WT, but would not be changed in Pgr-KO.

### Comparison of genes that are important for ovulation in human, mouse and zebrafish

Human (E-MTAB-2203) and mouse (GSE4260) [1, 3] transcriptomic data sets, that were focused on differentially regulated genes in the follicular cells of preovulatory oocytes were downloaded from EMBL and GEO databases, respectively. Two important marker genes, LHCGR and PGR, were both found in these two data sets. Therefore, both human and mice samples contain granulosa cells, and are comparable to the follicular cells of zebrafish.

Multiple CEL data files that containing human transcripts were first imported into Expression Console software (Affymetrix) for data normalization, then, the robust Multi-array Average (RMA) normalized our data and were transferred into Transcriptome Analysis Console program (v3.0, Affymetrix) for differential expression analysis using One-Way Repeated Measure ANOVA (paired). We used the R packages Affy [14] and limma [15] for differential expression analysis, due to only two biological replicates in mice samples.

Differentially expressed genes are defined as having a minimal 2 fold difference in the expression of the transcript observed in treated or mutant samples, compared to controls (absolute log2FoldChange > 1) with a corrected FDR p-value < 0.05. Ensembl gene IDs of human and zebrafish were converted to the mouse version to determine ovulatory genes that are conserved between human, mice, and zebrafish.

### Enrichment analysis

The Enrichment Map plugin for Cytoscape [16] was used to visualize the results of enriched gene sets and pathways due to Pgr-KO. Both up-regulated and down-regulated genes were first imported into the online g:Profiler (http://biit.cs.ut.ee/gprofiler/). We selected the “no filtering” option and kept gene sets that had three or more genes that were significantly differentially expressed for follow-up analyses. These gene sets were further compiled using KEGG, Reactome, and GO databases. Enriched gene sets were then loaded into the Enrichment Map plugin for Cytoscape using a p-value cutoff at 0.0005 to achieve an operable clustered network. The resulting network map was curated manually by removing uninformative gene sets and grouping functionally related gene sets together. We then labeled these functional groups to highlight prevalent biological functions that were enriched.

### Statistical analysis

Differences in expression between Pgr-KO and WT data were analyzed using Students’ t-test, with p-values < 0.05 considered to be significant.

## Results

### A large set of genes (>10%) are differentially regulated prior to ovulation

Global gene expression in Pgr-KOs showed a significant difference and clustered distinctly from those in WT (Figs. 2 & 3). Variation in gene expression among three different Pgr-KO fish were much smaller than those in WT (Figs. 2 & 3), indicating that Pgr is a key transcriptional regulator, and knocking out Pgr blocks the changes of gene expression critical for ovulation. Variable increases between WT samples suggest that gene expression changed dramatically prior to ovulation, or small sample size.

**Figure 2.**
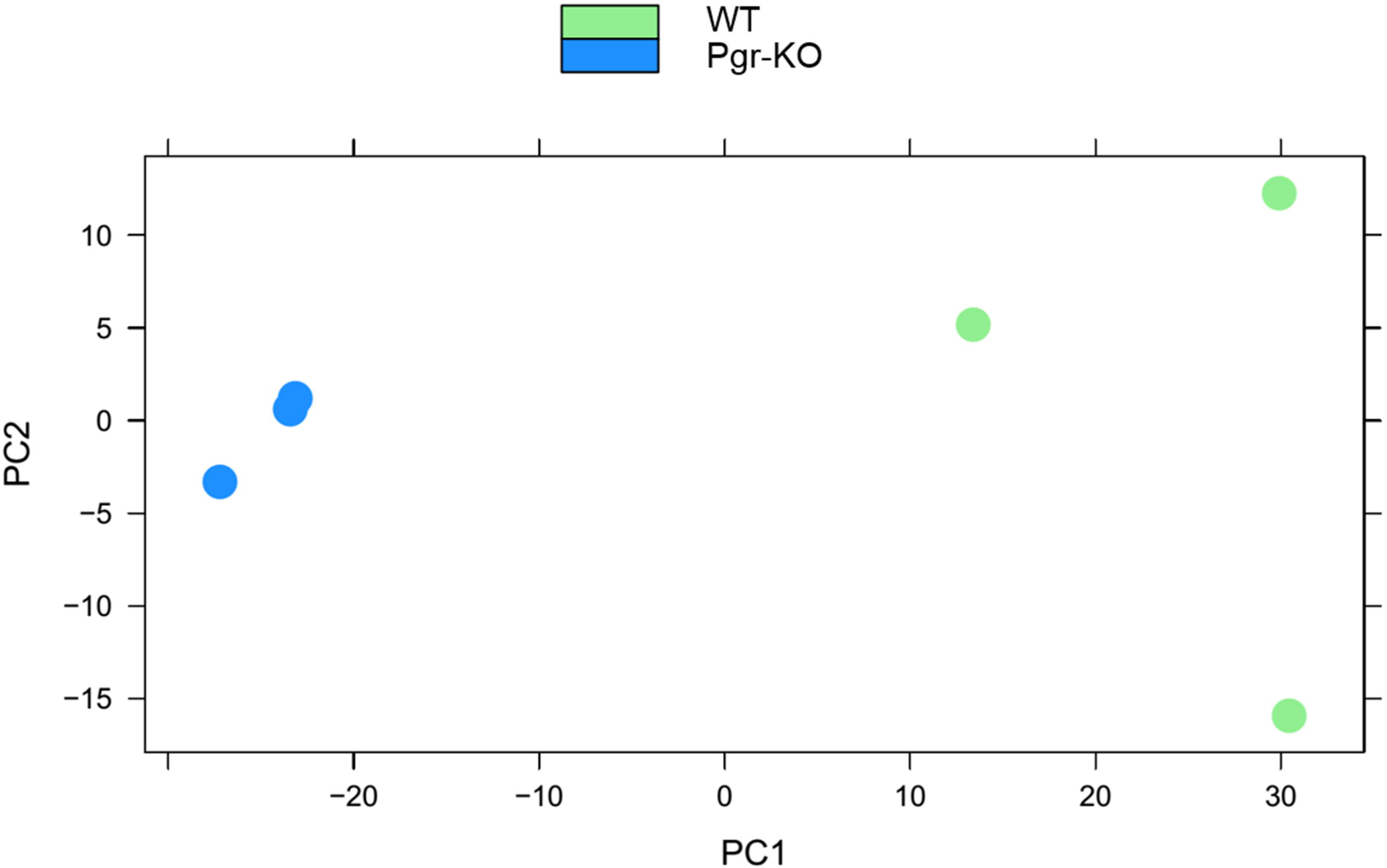
Principal component analysis (PCA) of transcriptomic data from pre-ovulatory follicular cells of Pgr-KO and wildtype (WT) zebrafish. The blue circles represent transcriptomic data from three pre-ovulatory follicular samples collected from three Pgr-KO fish, and the green circles represent the data from three samples of three WT fish.

**Figure 3.**
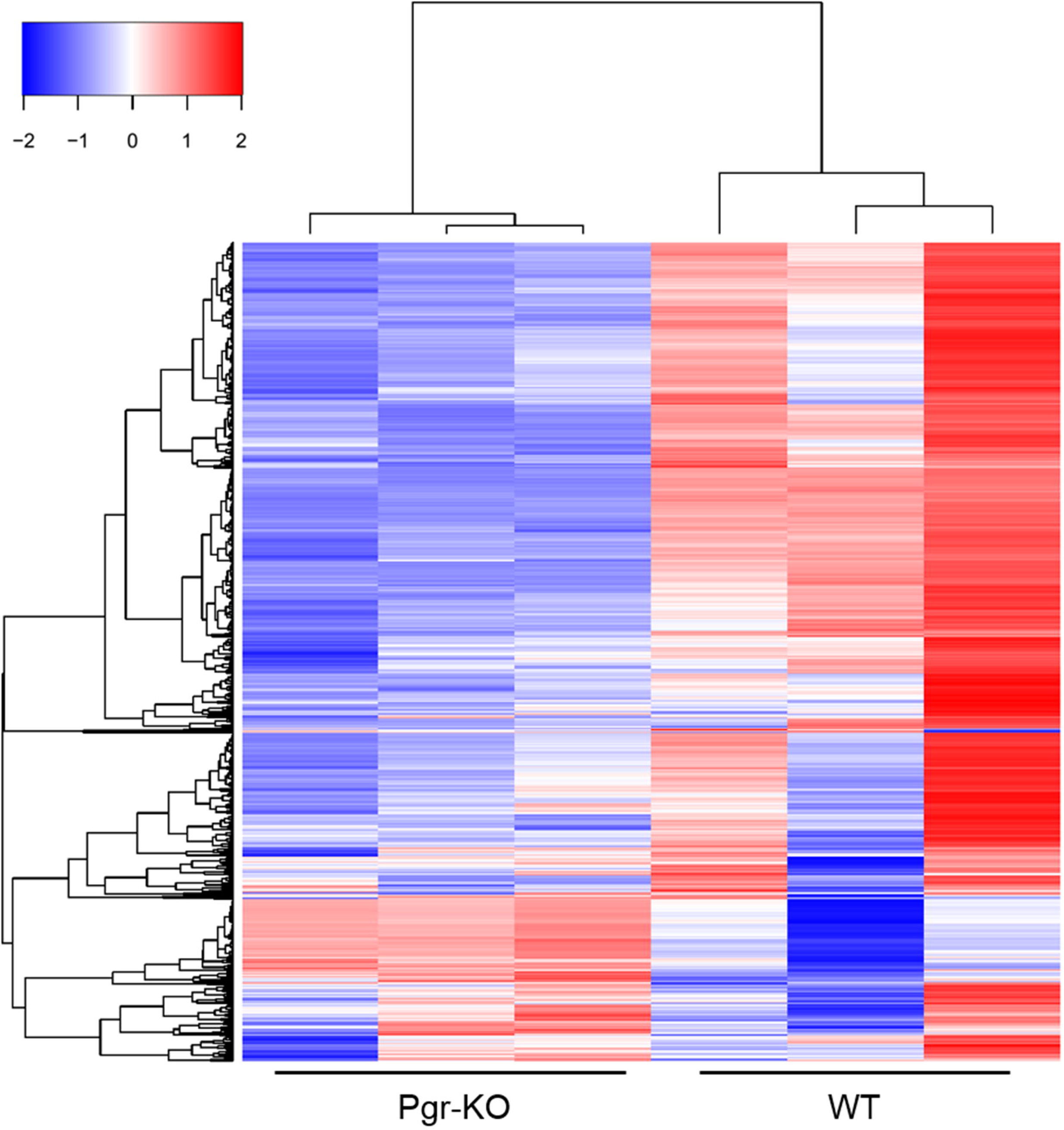
Heat map of top 2000 differentially expressed genes ranked by FDR-corrected p-value between WT and Pgr-KO. Each column represents an independent sample collected from follicular cells of stage IV oocytes that have completed final oocyte maturation, but prior to ovulation.

We found that 3,569 zebrafish genes are expressed significantly different in WT compared to those in Pgr-KO, using a cutoff of fold changes ≥ 2 and adjusted p-values≤0.05 (Table S2). We retained 2,888 significantly regulated zebrafish genes that have mouse homologs for subsequent analyses; and removed duplicates with less significance (Table S3). We did this to reduce the complexity caused by teleost specific gene duplication, and to search for conserved genes in vertebrates. Among these differentially regulated genes, 1,230 genes had higher expression and 1,658 genes had lower expression in WT than in Pgr-KO. Intriguingly, among the top 200 differentially expressed genes ranked by their adjusted p-value, 178 (89%) had significantly higher levels of expression in WT than in Pgr-KO (Fig. 4). They also exhibited large differences in expression, with 154 genes ranging eight to 147 fold. In contrast, only 16 genes among the top 200 had more than an eight fold decrease in WT than in Pgr-KO (TABLE S4). These results suggest dramatic changes in gene expression, especially increase of expression was critical for ovulation to proceed, which was also likely cause of variations among WT samples.

**Figure 4.**
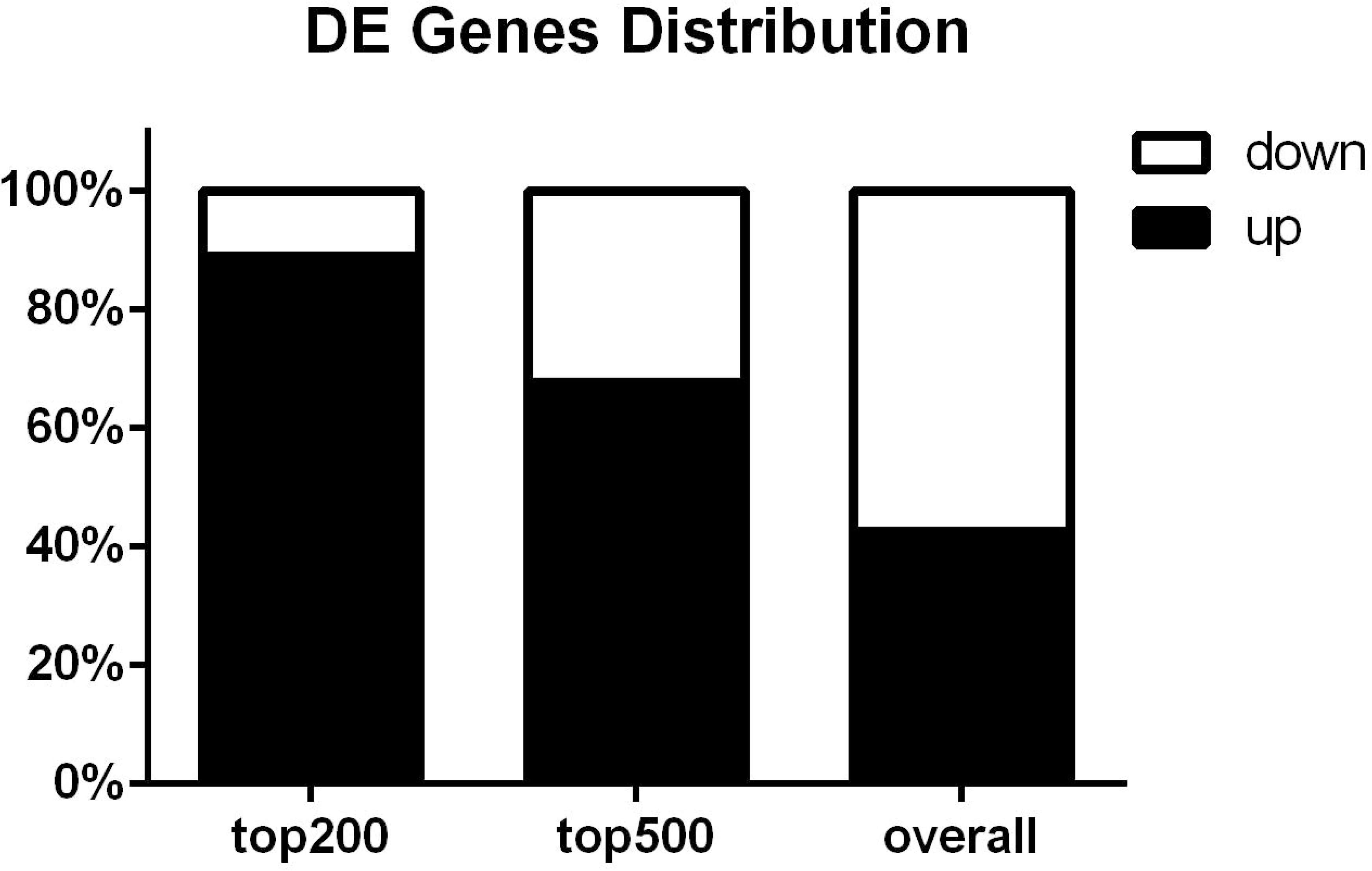
Distribution of up- or down-regulated genes in wildtype (WT) compared to Pgr knockout (Pgr-KO). Majority of top 500 genes are up-regulated in WT indicate increase of gene expression is important for ovulation.

### RNA-Seq results confirmed by qRT-PCR

The specificities of each PCR primer set used in qPCR was confirmed using a melting curve analysis and by sequencing the PCR products. Similar differences were obtained in the gene expression for 23 selected genes in qPCR analysis and RNA-Seq of WT samples compared to Pgr-KO (Fig.5).

**Figure 5.**
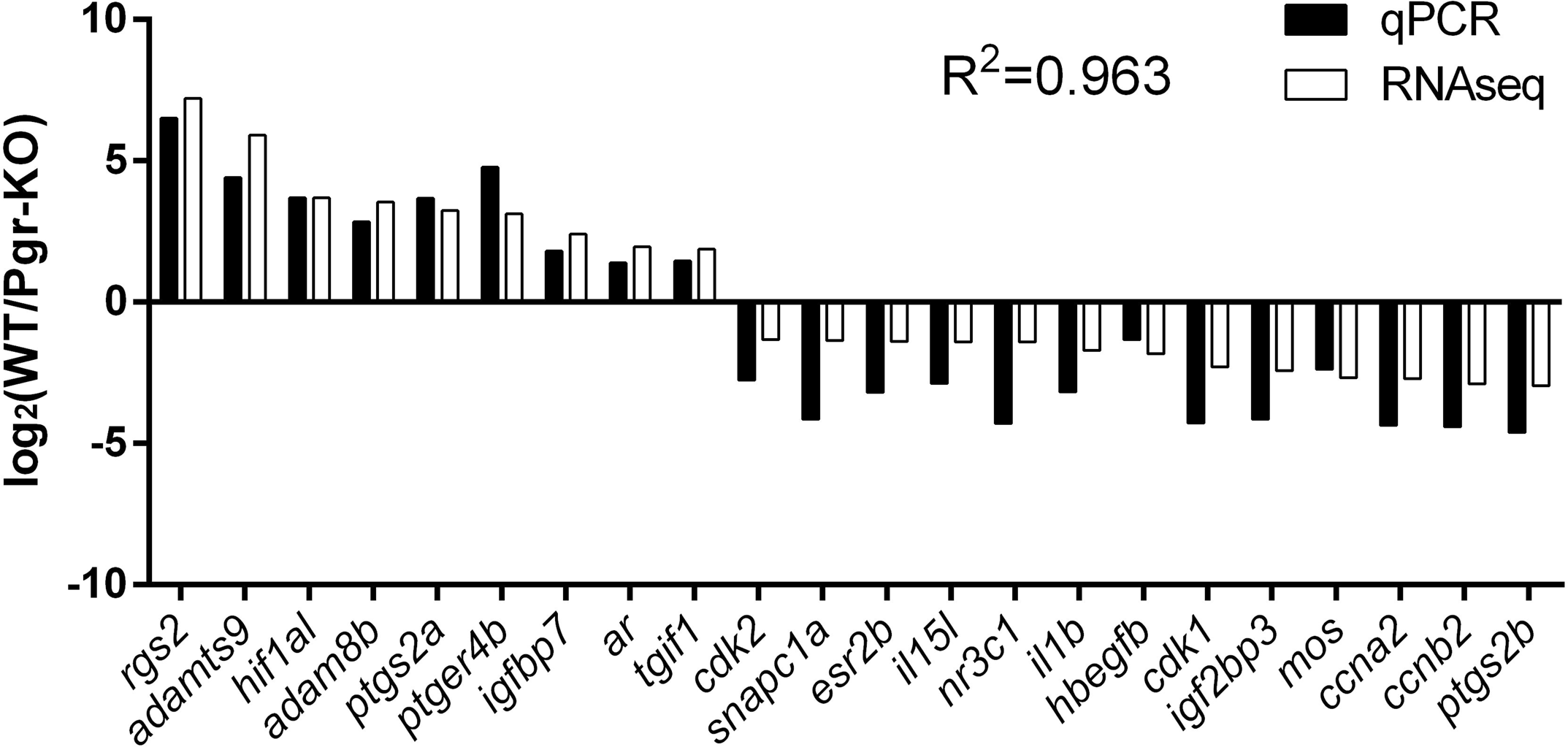
Validation of RNA-Seq results using real-time quantitative PCR (qPCR) based on relative fold changes of 23 selected genes in WT compared to Pgr-KO. Relative fold changes were expressed as log 2 of normalized average count (RNA-seq), or mean copy number (qPCR) of each gene in three WT samples (n = 3) divided by those respective numbers in three Pgr-KO (n = 3). The expression of *adamts1* was undetectable in the follicular cells by RNA-Seq and qPCR, and therefore was not included. Correlation coefficients (R^2^) between qRT-PCR and RNA-Seq based expression profiles is 0.963.

### Pgr is essential for regulating downstream targets that are important for ovulation

In total, 1,962 gene sets were enriched significantly (*p*≤0.05) when all up-regulated genes, i.e., 1,230 corresponding mouse homolog IDs in WT fish were input into g:Profiler. A small 943 gene sets were enriched when all down-regulated genes, i.e., 1,658 mouse homolog IDs, were input into the same program. We retained 649 upregulated gene sets and 469 down-regulated gene sets, with a more stringent cutoff at p≤0.0005, to draw better and less congested enrichment maps.

Multiple sets of genes, signaling pathways, and biological processes important for ovulation had significantly higher expression in WT compared to those in Pgr-KO. These enriched biological processes in WT include: angiogenesis, cell migration, chemotaxis, focal adhesion, response to growth factor, vasodilation, blood coagulation, cytokine production, inflammatory response, leukocyte aggregation and differentiation, cytoskeleton reorganization, extracellular matrix organization, response to hypoxia, and apoptosis (Fig.6A). The enriched KEGG or Reactome pathways in WT include: MAPK signaling, Wnt signaling, ERBB signaling, PI3K-Akt signaling, and NF-kappaB signaling (Fig.6A). In contrast, biological processes enriched in the WT down-regulated gene sets were mainly related to cell growth, proliferation, and cell cycle instead of ovulation [3]. These processes include: DNA replication, DNA repair, DNA methylation, cell phase transition, and cell division (Fig.6B). The clear distinction between up- and down-regulated genes in WT compared to Pgr-KO demonstrates that Pgr is essential for determining the cell fate and transition of pre-ovulatory follicular cells. A lack of Pgr would block the biological processes and signaling pathways required for ovulation.

**Figure 6.**
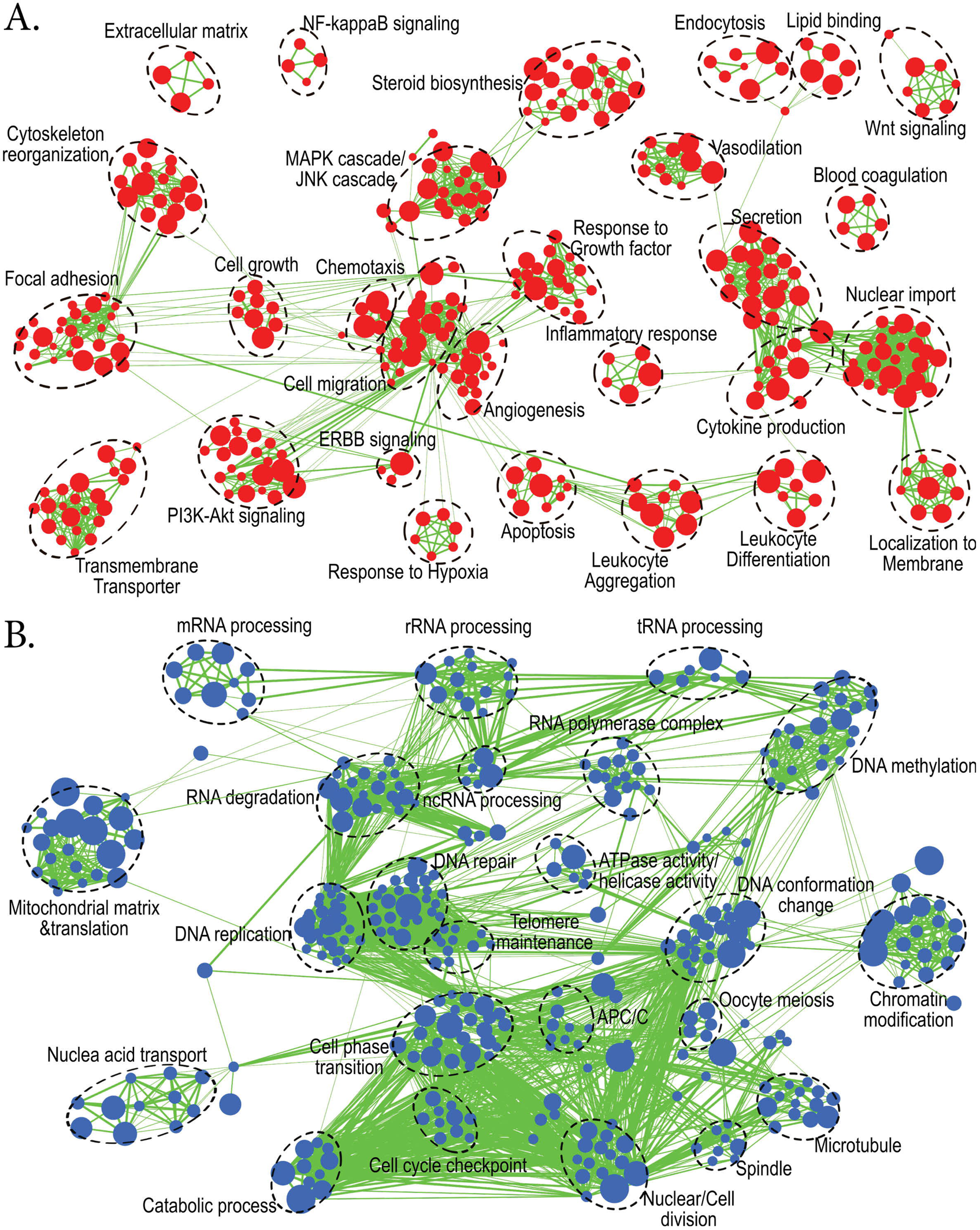
Enrichment map analysis showed two groups of biological processes and pathways, which were significantly enhanced or suppressed in wildtype (WT) compared to Pgr-KO (p = 0.0005). Nodes represent enriched gene sets, clustered automatically by the Enrichment Map plugin for Cytoscape program [16] according to the number of genes shared within sets. Node size is proportional to the total number of genes within each gene set. Proportion of shared genes between gene sets are represented by the thickness of the line between nodes. Functionally related gene sets were manually grouped, encircled by a dashed black line, and labeled. **A**. Enrichment map for 1230 up-regulated genes in WT compared to Pgr-KO. **B**. Enrichment map for 1658 down-regulated genes in WT compared to those in Pgr-KO.

### Conserved biological processes and pathways that are important for ovulation in human, mouse and zebrafish

Currently, Pgr-KO models are limited. Other than zebrafish, there is only one data file, containing no-biological replicates for a PGR-KO mouse and used whole ovaries, which could not be analyzed appropriately [17]. In order to further examine conserved genes and biological processes important for ovulation, we compared our RNA-Seq data with transcriptomic data obtained from HCG induced ovulation samples in humans [3] and mice [1]. In follicular cell samples of human treated with HCG, 852 genes were significantly up-regulated and 884 were down-regulated when compared to those without HCG treatment. Whereas, in mouse follicular cell samples treated with HCG, 1,356 genes were significantly up-regulated and 1,553 were significantly down-regulated compared to those without HCG exposure. We focused on analyzing genes that showed similar increases or decreases between zebrafish and mammals (human and/or mouse), and found 283 up-regulated genes and 378 down-regulated genes in WT that are conserved (Fig.7). The features of these three datasets are summarized in Table 1.

**Figure 7.**
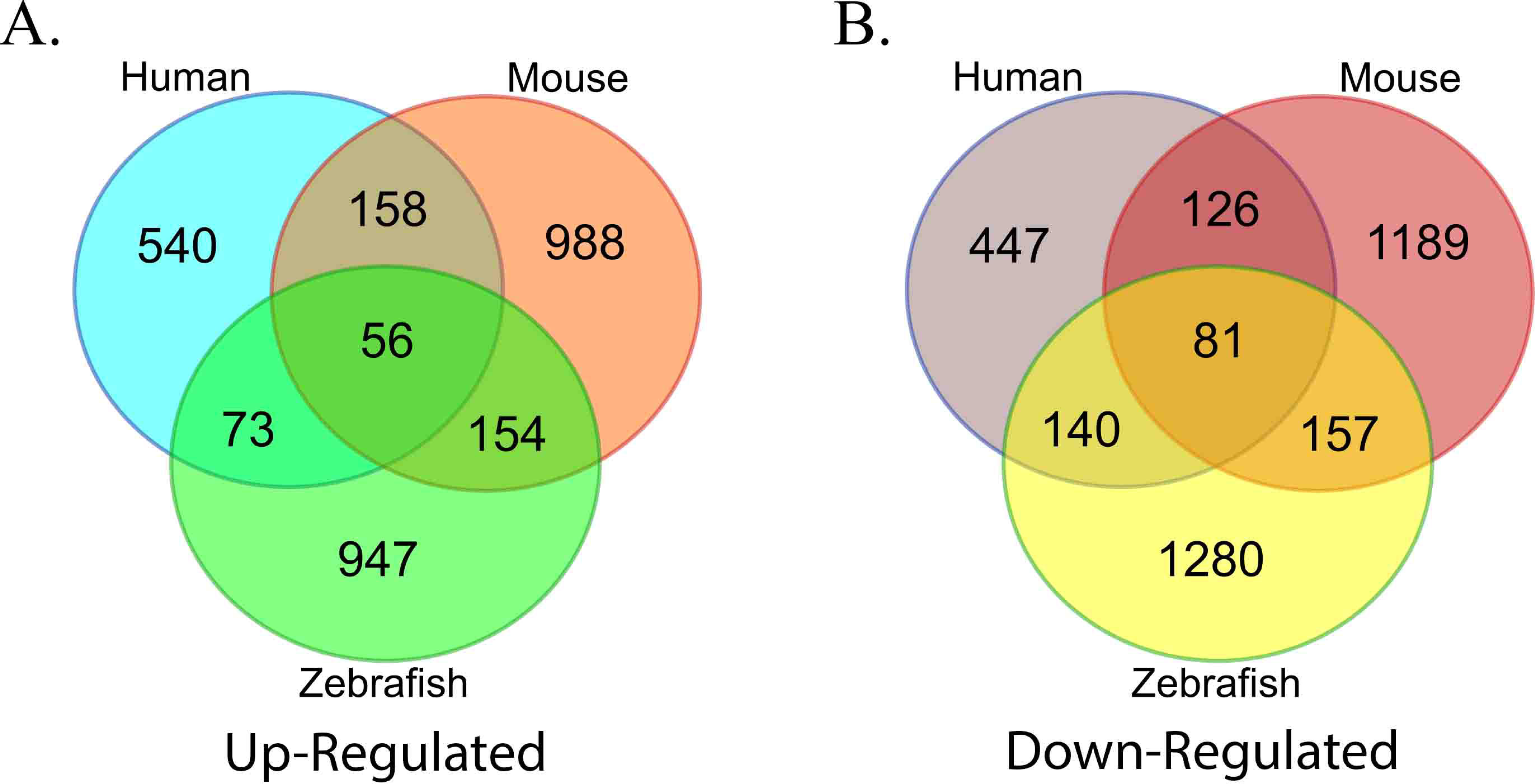
Comparison of significantly regulated genes in the follicular cells of zebrafish, human, and mouse during ovulation. Ensembl numbers of genes in each group that were derived from differentially regulated genes in human or zebrafish were converted to their respective mouse homologs. A. Up-regulated genes in WT zebrafish that underwent natural spawning cycle likely exposed to LH in vivo, in human or mouse follicular cells treated with HCG. A total of 283 genes (73 + 56 + 154) were up-regulated in WT fish during ovulation, as well as in humans and/or mice. B. A total of 378 genes (140 + 81 + 157) were down-regulated in WT zebrafish, humans or mice during ovulation.

**Table 1.**
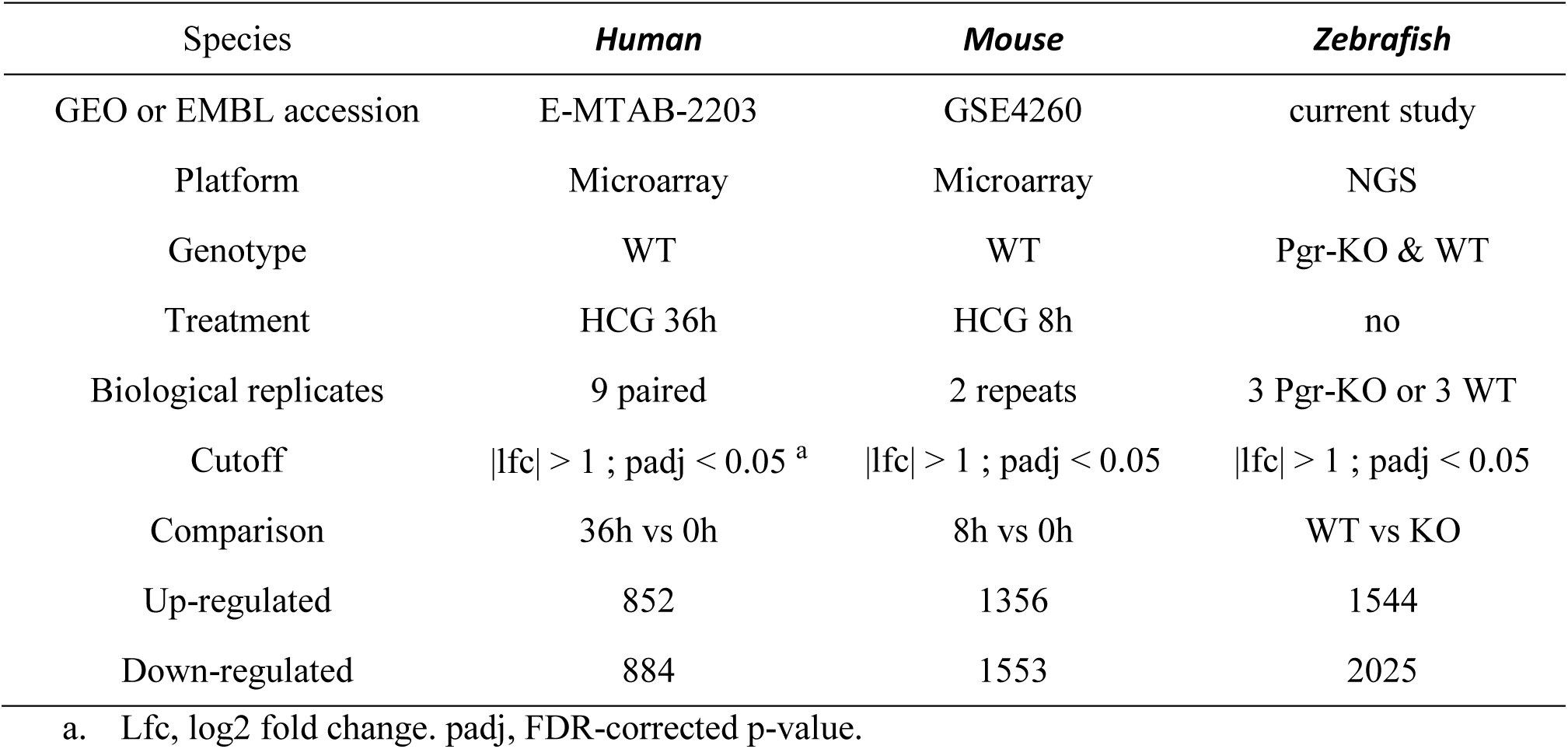
Comparison of three transcriptomic experiments for analyzing differentially regulated genes during ovulation in the follicular cells of zebrafish, human and mouse.

Enrichment analysis of the conserved 283 up-regulated or 378 down-regulated genes show two distinct pathways (Fig.6). The first pathway was related to ovulation including angiogenesis, cell migration, cytokine production, inflammatory response, MAPK/JNK cascade, and cell-specific apoptosis. The second was related to cell growth and proliferation, including DNA repair, DNA replication, chromatin modification, nuclear division, and cell cycle checkpoint (Fig.8). Our analysis indicates that these genes and biological processes for ovulation are highly conserved among vertebrates.

**Figure 8.**
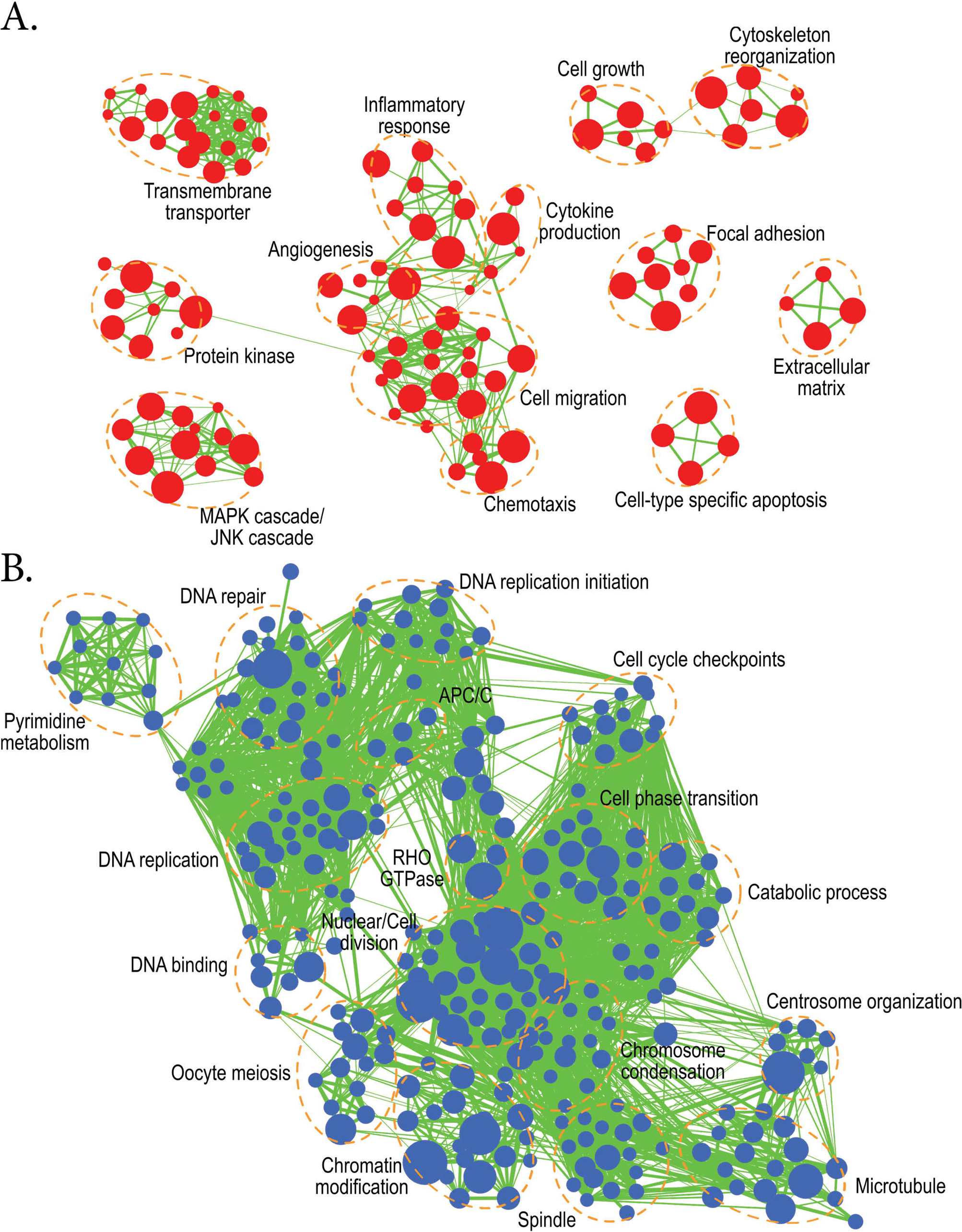
Enrichment map analysis using conserved 283 up-regulated and 378 down-regulated genes shared between fish and mammals (human and/or mice) during ovulation (p = 0.0005). **A**. Up-regulated biological processes during ovulation which include: transmembrane transportation, protein kinases, MAPK/JNK cascades, angiogenesis, the inflammatory response, cytokine production, cell migration, chemotaxis, extracellular matrix organization, cell growth, and cytoskeleton reorganization. **B**. Down-regulated biological processes during ovulation include: Pyrimidine metabolism, DNA repair, DNA methylation, DNA binding, DNA replication, cell cycle checkpoints, oocyte meiosis, chromatin modification, and cell phase transition.

### Representative genes and processes that may be important for ovulation

Some of the genes and processes conserved in our cross-species comparison (Fig.8), or suggested to be important for ovulation are highlighted below (Table. 2).

**Table 2.**
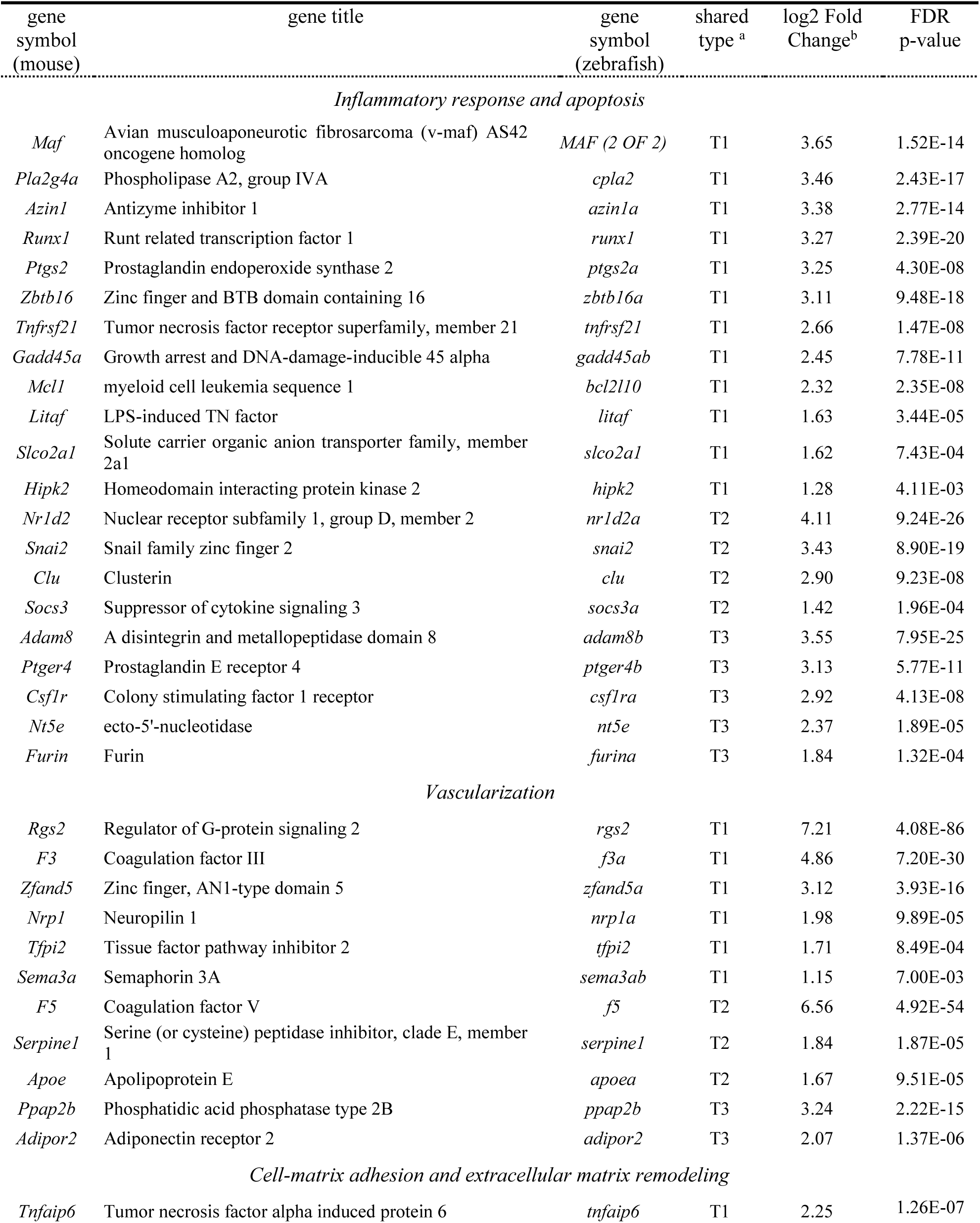
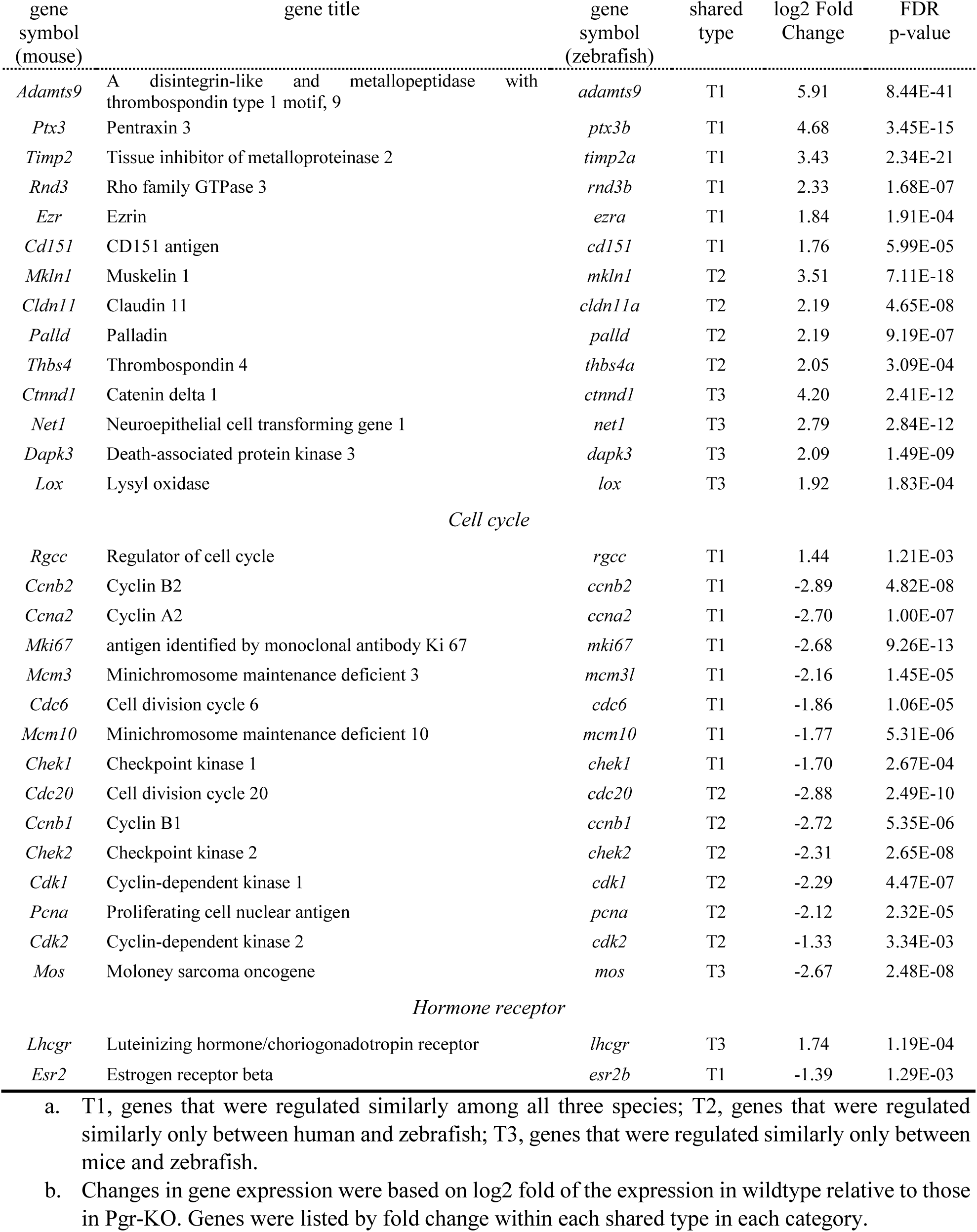
Representative differentially regulated genes in zebrafish during ovulation that are conserved between zebrafish, human and mouse.

#### Inflammatory response

Several inflammation-associated gene transcripts were more expressed in WT compared to those in Pgr-KO zebrafish, and higher in mice and human samples treated with HCG. Several of these genes are critical for ovulation. For example, *runx1* (runt related transcription factor 1), *ptgs2* (prostaglandin-endoperoxide synthase 2), and *pla2g4a* (phospholipase A2 group IVA) are important for prostaglandin synthesis and ovulation. Genes involved in il6 signaling include *socs3* (suppressor of cytokine signaling 3), and *mcl1* (myeloid cell leukemia sequence 1), which were also expressed significantly higher in WT compared to Pgr-KO fish. *Zbtb16* (zinc finger and BTB domain containing 16), *adam8* (a disintegrin and metallopeptidase domain 8) and *ptger4* (prostaglandin E receptor 4), which were also expressed significantly higher in WT compared to Pgr-KO. Intriguingly, *tnfrsf21* (tumor necrosis factor receptor superfamily, member 21) and *furin* (paired basic amino acid cleaving enzyme) transcripts were also expressed more in WT than Pgr-KO.

#### Vascularization

Genes involved in coagulation and angiogenesis including *F3*, *F5* (coagulation factors F3 and F5), and plasminogen activator inhibitor *serpine1* (serine peptidase inhibitor clade E member 1) were expressed significantly higher in WT compared to those in Pgr-KO. Notably, the expression levels of *rgs2* (regulator of G-protein signaling 2) encoding a member of GTPase activating proteins, and *nrp1* (neuropilin 1) encoding a protein that has been shown to interact with VEGFA (vascular endothelial growth factor A), were both expressed significantly higher in all three species prior to ovulation in WT.

#### Extracellular matrix remodeling

Extracellular matrix disassociation is an essential step for follicle rupture. Surprisingly, transcripts of *adamts1* (a disintegrin-like and metallopeptidase with thrombospondin type 1 motif 1), suggested to play a key role in mammalian COC expansion leading to follicular rupture [18], was undetectable in zebrafish follicular cells in both RNA-Seq and qPCR analyses though *adamts1* was highly expressed in other cell types including follicles in earlier stages (unpublished). Instead, another member of Adamts family, *adamts9,* was found to be up-regulated 60 fold in WT samples compared to Pgr-KO.

#### Cell cycle

Most of cell cycle related genes (14 out 19) were up-regulated in Pgr-KO, but during onset of ovulation in WTs were significantly down-regulated. Similar trends of decrease or increase were observed in all three species. These genes include several cyclins (*ccnb2*, *ccna2,* and *ccnb1*), cyclin-dependent kinases (*cdk1* and *cdk2*), a cell proliferation marker (*mki67*), several mini-chromosome maintenance proteins (*mcm3*, *mcm4*, *mcm6*, *mcm7,* and *mcm10*), related to cell division cycle (*cdc20*), and cell phase transition (*mos*).

#### Steroid and hormone receptors

Compared to Pgr-KO, the expression of *esr2* (estrogen receptor beta) was lower in WT. Similarly, the expression of *Esr2* was lower in human and mice samples treated with HCG. On the other hand, the expression of *lhcgr* (Luteinizing hormone/choriogonadotropin receptor) was significantly higher in WT than Pgr-KO fish, with similar results observed in mice but not in human samples.

## Discussion

We have performed the first genome-wide differential gene expression analysis that is specifically designed for the follicular cells of pre-ovulatory oocytes in the zebrafish, and conducted the first comparison of differentially regulated genes in the follicular cells of pre-ovulatory oocytes among zebrafish, mice, and humans. Our analysis indicates that ovulation is involved in a large network of approximately 3,000 genes, which is 11.5% of 26,000 available genes in zebrafish, all working in concert in the follicular cells of pre-ovulatory zebrafish oocytes. The number of ovulation related genes found in the zebrafish is about three times more than those reported in humans, but only slightly more than those reported in mice (Table. 1). One likely reason is that RNA-Seq is much more sensitive than microarrays used in previous studies. Or, it is possible that we overestimate the ovulation-related genes in zebrafish due to differences in gene expression that may be present before ovulation occurs in Pgr knockout. It is impossible to predict which WT fish will undergo ovulation unless they have completed final oocyte maturation (see Fig.1 for detail). Not all fish ovulate daily, so we did not collect follicular cells prior to maturation in current study due to lacking a reliable marker to identify which fish will ovulate before the completion of oocyte maturation in vivo. Therefore, it will be necessary to establish an ovulation assay in vitro and to study gene expression during ovulation. Nevertheless, by comparing our data sets with mammalian data sets we are able to show many conserved genes are related to ovulation and associated signaling pathways are very similar whether using the entire set of differentially regulated zebrafish genes or conserved “ovulatory” genes of fish, mouse, and human. These differentially expressed genes activate signaling pathways and biological processes that are important for ovulation to precede while also down regulating signaling pathways involved in growth and proliferation prior to ovulation. The switch from a period of growth and proliferation to the ovulation is important for follicular cell rupture and the process to proceed appropriately. These ovulatory genes and signaling pathways are highly conserved among fish, mice and humans.

Therefore, as demonstrated herein and elsewhere, zebrafish offer an alternative model for studying molecular mechanisms and biological processes that control ovulation.

As we noted at beginning, biological processes leading to the ovulation are quite different between mammalian models and zebrafish model, mainly due to in vivo gestation in mammals in comparison to external fertilization and development in zebrafish. We were surprised to find so many conserved “ovulatory” genes as well as a significant number of non-conserved “ovulatory” genes (Fig.7). A large number of non-conserved “ovulatory” genes is likely due to difference in experimental treatments, different analytic tools (RNA-seq vs microarray), difference in cell population, or different molecular mechanisms in control of ovulation in different models. For accurate comparison, we have to generate transcriptomic data sets using same sensitive sequencing technique, and same genotype (PGR KO) in different model species. We will focus our discussion on conserved genes and pathways potentially important for ovulation, since more information is required for defining different ovulatory mechanisms in different species.

### Inflammatory response

Substantial evidence from studies in mammals has shown that biological events occurring in an ovulating follicle are similar to those in an acute inflammatory response [2, 3, 19]. One major indicator is the increase of prostaglandins (PGs) resulting from the upregulation of Ptgs2, a rate-limiting step in prostaglandin biosynthesis. It is still unclear whether PGR and PTGS are two separate downstream pathways of LH signal, as no evidence shows promoter binding sites for PGR on *Ptgs2*. Some have suggested that several genes including Pparg and Il6 may act as mediators between PGR and PTGS2, as demonstrated by conditional Pparg knockout mice [19]. However, the transcripts of these mediators were not significantly regulated in the follicular cells of pre-ovulatory oocytes in zebrafish, implying Ppar pathway might not be an important mediator in zebrafish ovulation. Another transcriptional factor, RUNX1 is a candidate mediator since upregulation of *Ptgs2* by RUNX1 can be inhibited by a PGR antagonist, and direct binding of RUNX1 to two RUNX-binding motifs in the *Ptgs2* promoter region has been confirmed by CHIP and EMSA assays [20, 21]. Our finding of significantly upregulated *Runx1* (fold changes > 10, P < 2.40E-20) in the follicular cells of pre-ovulatory oocytes supports that RUNX1 as a mediator regulating *Ptgs2*. On the other hand, the present study shows one of the PGs receptors, *Ptger4,* was markedly up-regulated prior to ovulation in both mouse and zebrafish datasets. It has been shown that Pgr directly regulates *ptger4* in the pre-ovulatory follicles of medaka [7, 22]. In contrast, the role of PTGER4 in ovulation has been overlooked in mammals as studies focused more on role of PTGER2 in COC expansion and ovulation [23–25]. Our studies provide possible candidates that involve in PGR signaling pathway and ovulation. Clearly, more studies are needed to understand the molecular mechanisms that regulate members of PTGS and PTGER and their functions in ovulation.

### Vascularization

Vasodilation induced by LH in the pre-ovulatory ovary is required for increased vascular permeability. This drives leukocytes to migrate from the blood vessels to the interior of the pre-ovulatory follicles to release multiple cytokines and elicit inflammatory reactions leading to follicle wall breakdown [19]. The expression of *rgs2*, which shows the greatest increase during ovulation in our RNA-Seq data (>100 fold), has been suggested to be essential for stabilizing blood pressure via inhibition of Gq/11-mediated signaling in human cardiovascular system [26, 27]. Evidence also suggests that *Rgs2* expression can be stimulated by LH, but is attenuated by PGR antagonist or PTGS2 inhibitor in the pre-ovulatory follicles of mice and bovine [28, 29]. It remains to be elucidated whether or not RGS2 can play a similarrole in follicle rupture as it does in cardiovascular system, increasing vascular permeability.

NRP1, a well-known membrane bound co-receptor that is involved in VEGF signaling and vascularization had increased expression in pre-ovulatory follicles of zebrafish, mouse, and human. NRP1 up-regulates KDR downstream signaling in response to VEGF in angiogenic modulation [30]. We also found a significant increase of *kdr* expression in the follicular cells of pre-ovulatory oocytes in zebrafish. To prevent microbleeding during ovulation, multiple coagulation factors such as *f3*, *f5,* and tissue factor pathway inhibitor 2 (*tfpi2*) are concomitantly up-regulated in zebrafish, consistent with previous findings in humans [3]. The up-regulation of *serpine 1(pai1),* a blood clotting promoting factor, before ovulation in zebrafish is consistent with the recent finding in medaka. Serpine1 has been suggested to be a limiting regulator controlling plasmin hydrolyzing laminin, a major basement membrane component situated between the granulosa and theca cells of the follicle in medaka [31]. However, since PAI1 deficient mice are viable, the role of PAI1 in vascularization during ovulation should be further explored [32, 33].

### Extracellular matrix remodeling and follicle wall breakdown

Proteases are required for extracellular matrix degradation and remodeling, and are essential for follicular cell rupture and the release of mature oocytes. The involvement of several metalloproteinases including members of MMP (matrix metalloproteinase), ADAM, and ADAMTS families have been examined [34–36]. However, evidence of these enzymes being involved in ovulation is limited, partly due to lack of obvious effects on ovulation or embryonic lethality in knockouts [37]. Interestingly, expression and knockout studies of *Adamts1* have suggested the involvement of ADAMTS1 in ovulation and fertility in mammals. *Adamts1* was up-regulated in the pre-ovulatory follicles of several mammalian species. This up-regulation was partly dependent on the expression of *Pgr* in the granulosa cells. Knockout of *Adamts1* in mice suggests an important role of this protease as a downstream effector of PGR in ovulation. Homozygous *Adamts1* knockouts are subfertile, producing litters 4- to 5-fold less than control littermates, partially due to failed rupture of some large follicles [36]. *Adamts1* knockout mice are subfertile and have less severe phenotypes than *Pgr* knockout mice, who cannot ovulate and therefore have no litters. This indicates that ADAMTS1 may not be a key protease, or other PGR regulated proteases contribute to the ovulatory mechanism. Lacking significant differences in gene expression, enzyme concentration, or enzymatic activity of ADAMTS1 in infertile women compared to controls also did not support functional roles of ADAMTS1 in human fertility [38]. ADAMTS1 has been suggested to cleave versican in COC matrix [37, 39], so the functions of this gene in basal vertebrates may be different since no COC is necessary.

Intriguingly, upregulation of *adamts9* in mammals and zebrafish during ovulation suggests that this enzyme is most likely involved in ovulation, and its function conserved across vertebrate species. Adamts9 was found to be upregulated in pre-ovulatory follicles or GCs following HCG treatment in macaque and human [2, 3, 40]. GON-1, an ortholog of ADAMTS9, is involved in the degradation of extracellular matrix (ECM) and is essential for gonadal morphogenesis in *C. elegans*. GON-1 helps migration of distal tip cells by degrading extracellular matrix components. In GON-1 mutants, the adult gonad is severely disorganized with no arm extension and no recognizable somatic structure. The developmental defects in gon-1 mutants are limited to the gonad; other cells, tissues, and organs develop normally in *C. elegans*. However, functions of ADAMTS9 in vertebrates have not been established, partly due to *Adamts9* knockout mice dying before gastrulation [41]. Thus, using alternative models or establishing conditional knockouts are required for examining the function of ADAMTS9 in vertebrates.

### Missing information was compensated by our fish data

Some important ovulatory genes and signaling pathways such as ERBB, PI3K, and WNT signaling are unexpectedly missing in the shared gene lists and conserved enrichment maps, but significantly enriched in zebrafish data set (Table 3; Fig. 6). ERBB signaling (epidermal growth factor receptor signaling) is induced by the LH surge through the activation of EGF-like factors [42]. The increased expressions of *lhcgr*, *egfr,* and *mapk1* along with downstream genes (e.g. *ptgs2*, *tnfaip6*) in the follicular cells of WT pre-ovulatory zebrafish oocytes suggest that ERBB signaling is conserved and involved in the ovulation of vertebrates. Knockouts of PTEN inhibited PI3K signaling, increased susceptibility to apoptosis and enhanced ovulation in mice [43]. We also found that *pten* (a tumor suppressor) was highly expressed in the follicular cells of WT, and PI3K/Akt signaling was enriched in up-regulated genes (Fig.6). Similarly, tumor suppressor genes *dab2ip* and *foxo3* were expressed much higher in WT, which correlated to WNT signaling being enriched in zebrafish. FOXO3 has been shown to inhibit WNT signaling and cancer development [44, 45]. Notably, we found at least 232 human tumor suppressor [46] homolog genes that were differentially expressed in the follicular cells of zebrafish pre-ovulatory oocytes. Molecular mechanisms and functions of these tumor related genes during vertebrate ovulation is still unclear.

Hypoxia typically occurs at the time of ovulation [47]. This is supported by high expression levels of *hif1al* (hypoxia-inducible factor 1, alpha subunit like) in the follicular cells of WT pre-ovulatory oocytes. Hypoxia response also activates angiogenesis, consistent with the high expression of vascular endothelial growth factor receptors *flt1* and *kdr* in WT fish. Activation of NF-kB (nuclear factor kappaB) is a critical part of transcriptional response to hypoxia and the local inflammatory response [48, 49]. High expression of *rela/p65,* a component of NF-kB, increased expression of *nfkbia* (NF-KappaB inhibitor, alpha) encoding the cognate binding inhibitor of Rela, and several ubiquitin related genes (*ubb*, *ubc*, *uba52*, *ube2a,* Table 3). This suggests possible activation of NF-kB signaling in pre-ovulatory follicular cells in response to hypoxia-like conditions of ovulation [50]. The high expression of *tnfrsf21* (tumor necrosis factor receptor superfamily, member 21/death receptor 6), during ovulation suggests that activation of NF-kB signaling is also involved in local inflammation [51]. Intriguingly, *il1b* showed low expression in WT fish, although IL1B could induce direct binding of NF-kB to the promoter sequence of ADAMTS9 in human chondrocytes [52]. More studies are required to understand the activation and involvement of NF-kB in response to hypoxia, and its regulation of inflammation and ADAMTS9 during ovulation.

**Table 3.**
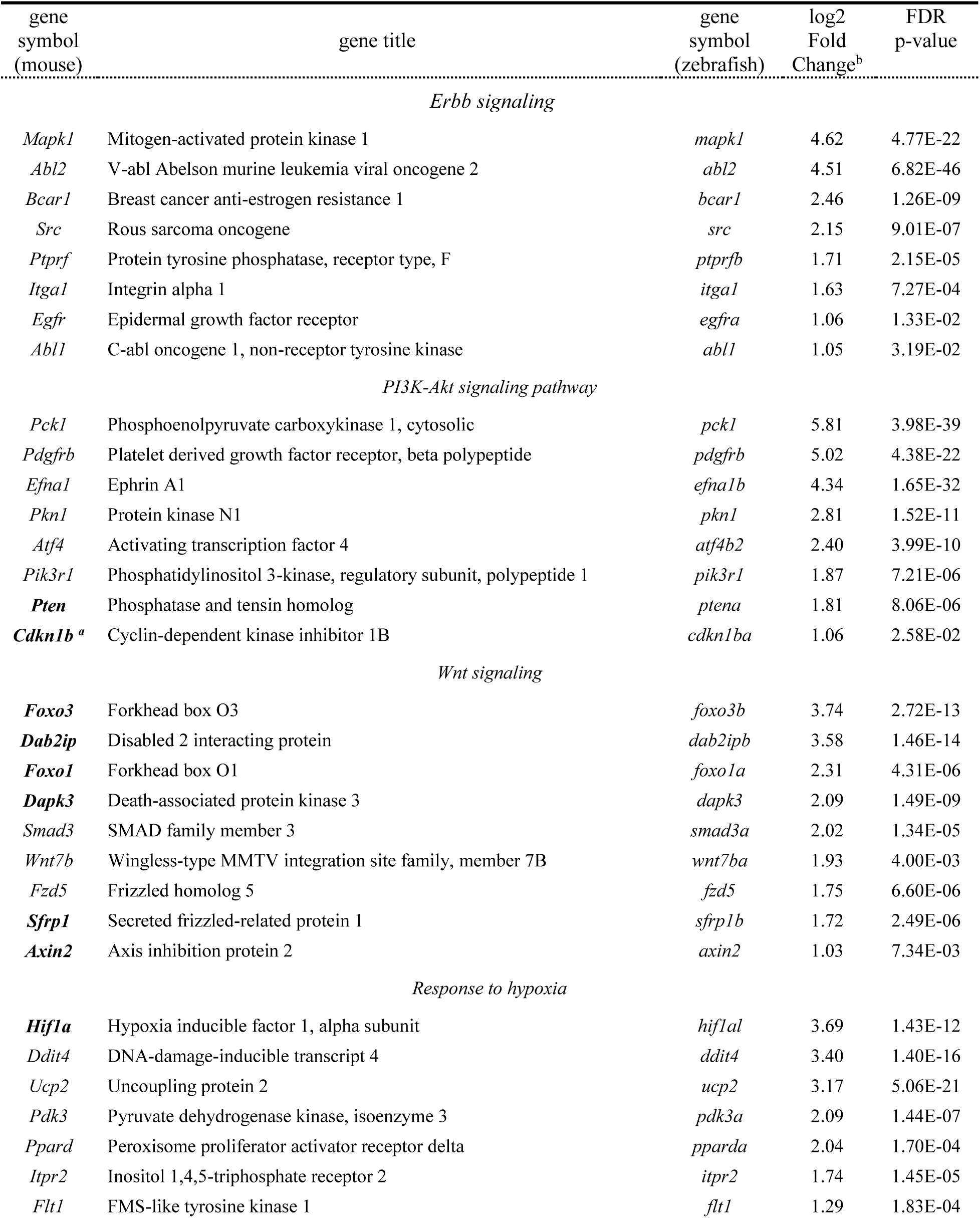
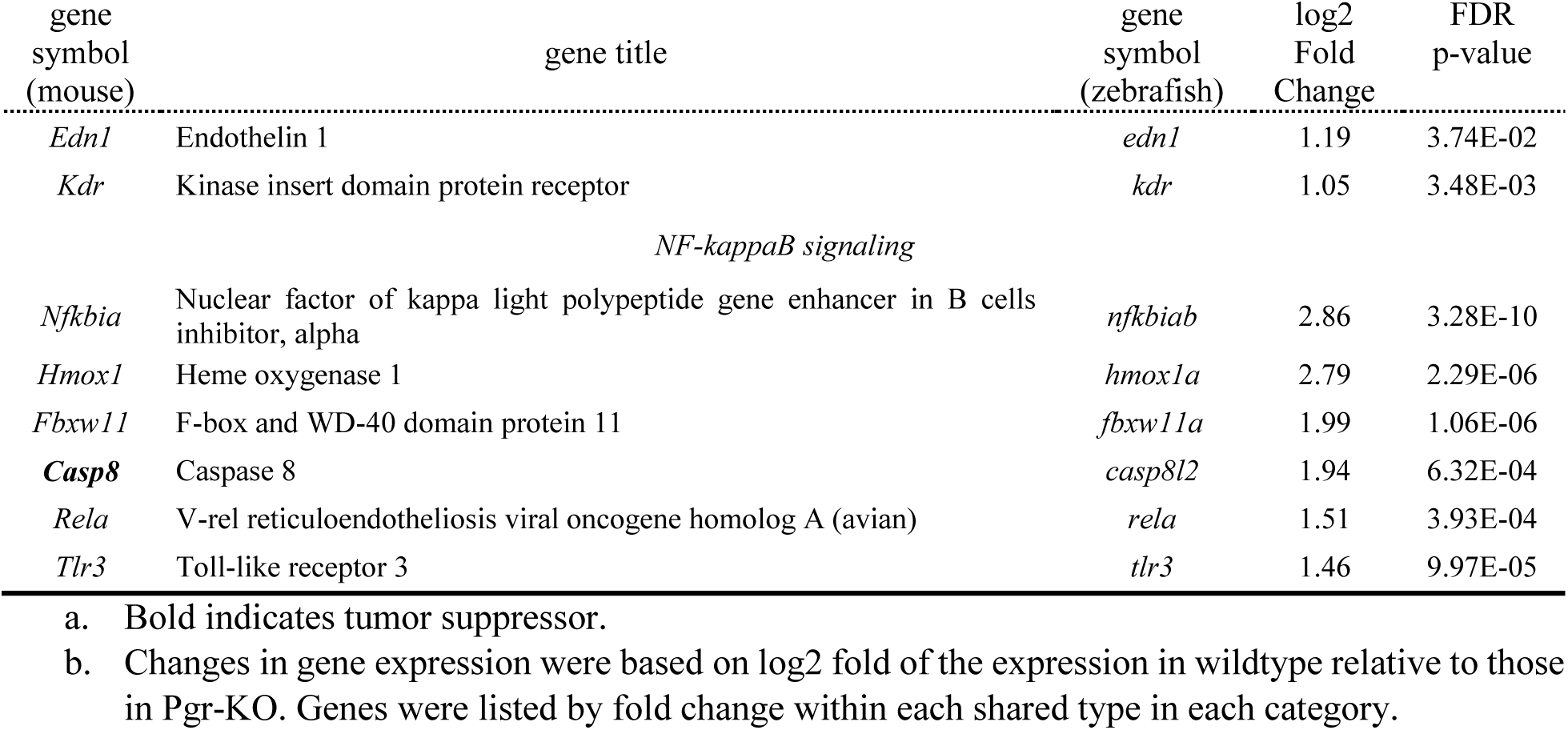
Representative genes and signaling pathways that are important for ovulation in zebrafish.

In summary, for the first time we successfully identified genes and signaling pathways that are potentially important for ovulation in zebrafish, a non-mammalian vertebrate model, using high-throughput sequencing and Pgr-KO. The comparison of differentially regulated genes among human, mouse, and zebrafish datasets further confirm that genes and signaling pathways important for ovulation are conserved among vertebrates. Zebrafish should serve as an excellent model for studying the function of genes and signaling pathways that are important for ovulation.

